# Genetic variation for mitochondrial function in the New Zealand freshwater snail *Potamopyrgus antipodarum*

**DOI:** 10.1101/087155

**Authors:** Joel Sharbrough, Jennifer L. Cruise, Megan Beetch, Nicole M. Enright, Maurine Neiman

## Abstract

The proteins responsible for mitochondrial function are encoded by two different genomes with distinct inheritance regimes, rendering rigorous inference of genotype-phenotype connections intractable for all but a few model systems. Asexual organisms provide a powerful means for addressing these challenges because offspring produced without recombination inherit both nuclear and mitochondrial genomes from a single parent. As such, these offspring inherit mitonuclear genotypes that are identical to the mitonuclear genotypes of their parents and siblings and different from those of other asexual lineages. Here, we compared mitochondrial function across distinct asexual lineages of *Potamopyrgus antipodarum,* a New Zealand freshwater snail model for understanding the evolutionary consequences of asexuality. Our analyses revealed substantial phenotypic variation across asexual lineages at three levels of biological organization: mitogenomic, organellar, and organismal. These data demonstrate that different asexual lineages have different mitochondrial function phenotypes and that there exists heritable variation (that is, the raw material for evolution) for mitochondrial function in *P. antipodarum.* The discovery of this variation combined with the methods developed here sets the stage to use *P. antipodarum* to study central evolutionary questions involving mitochondrial function, including whether mitochondrial mutation accumulation influences the maintenance of sexual reproduction in natural populations.

## INTRODUCTION

Mitochondrial function is of critical importance to eukaryotic health (for example,Chen *et al.,* 2007; Dowling, 2014), and genetic variation for mitochondrial function has been linked to evolutionary adaptation (for example,Rawson and Burton, 2002) and disease (DiMauro and Schon, 2001). The role of genetic variation for mitochondrial function is complicated by direct interaction between nuclear encoded and mitochondrially encoded proteins, particularly with respect to oxidative phosphorylation (OXPHOS) (reviewed in Rand *et al.,* 2004). Successful mitonuclear interaction is particularly important for proper enzyme function in OXPHOS complexes I, III, IV, and V because these complexes are composed of subunits encoded by both genomes. Accordingly, discordance between mitochondrial and nuclear genomes has been demonstrated to have negative fitness and/or functional consequences in a variety of animals, including copepods (Ellison and Burton, 2006), *Drosophila* (Meiklejohn *et al.,* 2013; Pichaud *et al.,* 2013), seed beetles (Dowling *et al.,* 2007), and salamanders (Lee-Yaw *et al.,* 2014).

In sexually reproducing organisms, the maintenance of mitonuclear compatibility is further complicated by the expectation that the different mechanisms of nuclear vs. mitochondrial genome (mtDNA) inheritance will differentially affect the generation, maintenance, and distribution of genetic variation. In particular, biparental inheritance and meiotic recombination in the nuclear genome should increase effective population size relative to the (typically) uniparentally inherited and non-recombinant mtDNA (reviewed in Barr *et al.,* 2005; Neiman and Taylor, 2009). This logic is the basis for the expectation that mtDNA will, when compared to the nuclear genome, experience reduced efficacy of selection and suffer an increased rate of accumulation of mildly deleterious mutations (reviewed in Neiman and Taylor, 2009). This mechanism is also thought to generate selection favoring compensatory changes in nuclear-encoded mitochondrial subunits (Sloan *et al.,* 2013; Zhang and Broughton, 2013).

Asexual taxa provide a particularly interesting context in which to evaluate mitochondrial function and evolution because the absence of recombination and segregation in asexually inherited nuclear genomes means that asexual lineages will transmit their mtDNA in complete linkage disequilibrium (LD) with the nuclear genome. To date, inbred sexual lineages have been the primary genetic tool used to investigate genotype-phenotype connections relating to mitochondrial function (for example,Ellison and Burton, 2006; Montooth *et al.,* 2010; Latorre-Pellicer *et al*., 2016). While these studies are powerful, the inferences that they generate are in part limited by the fact that inbreeding can introduce other off-target effects (for example, inbreeding depression, purging of harmful recessive mutations). Alternatively, full-factorial crossing of mtDNA onto various nuclear backgrounds (Willett and Burton, 2003; Dowling *et al*., 2007), perhaps represents the best of both worlds, but it is not tenable for species with long generation times. These challenges can be circumvented in asexual lineages, in which mitonuclear LD can be used as a relatively straightforward means of exploring genotype-phenotype connections in natural populations. This asexual-focused approach also has the substantial additional benefit of providing information relevant to understanding how sexual reproduction (and its absence) influences mitochondrial function.

*Potamopyrgus antipodarum,* a New Zealand freshwater snail, is very well suited for investigating mitochondrial function in the absence of sex because obligately sexual and obligately asexual individuals frequently coexist in natural populations (Lively 1987). Multiple separate transitions from sexual ancestor to asexual descendent have occurred within this species, providing many so-called “natural experiments” into the consequences of asexuality (Neiman *et al.,* 2011; Paczesniak *et al.,* 2013). Asexuality in *P. antipodarum* appears to occur via apomictic parthenogenesis (Phillips and Lambert, 1989), meaning that asexually produced offspring inherit both their nuclear and mtDNA from a single parent and that the nuclear genome is transmitted without recombination. The implications are that *P. antipodarum* individuals descended from the same mother (what we term “asexual lineages”) should share the same mitonuclear genotype, barring *de novo* mutations.

Here, we use a common-garden approach, which isolates genetic (*vs.* environmental) effects on phenotypic variation, to test whether distinct asexual lineages of *P. antipodarum* (our proxy for mitonuclear genotype) vary in mitochondrial function at three distinct levels of biological organization: (1) mitogenomic (mtDNA copy number), (2) organellar (mitochondrial membrane potential and electron transport), and (3) organismal (total oxygen (O_2_) consumption). All three of these traits have been linked to mitochondrial performance in other taxa. In particular, mtDNA copy number is thought to affect mitochondrial function (Van den Bogert *et al.,* 1993; Taanman *et al.,* 1997; Moraes, 2001; Salminen *et al.,* 2017) and to be dynamically regulated in response to various cellular environmental cues (Hori *et al.,* 2009; Matsushima *et al.,* 2010; Kelly *et al,* 2012). This regulation is also thought to be tuned, at least in part, as a response to the energy demands of a cell (Moraes, 2001). Elevated mtDNA copy number has even been shown to compensate for deletions in mtDNA (Bai and Wong, 2005), but whether copy number elevation represents a general compensatory mechanism for mitochondrial mutation accumulation remains unclear (Montier *et al,* 2009). Second, mitochondrial membrane potential, generated by electron transport, determines the strength of the electrochemical gradient mitochondria use to phosphorylate ADP to ATP (Nicholls, 2004). Variation in mitochondrial membrane potential has been linked to cellular aging (Nicholls, 2004) and longevity (Callegari *et al.,* 2011). The JC-1 assay measures the strength of the electrochemical gradient in mitochondrial isolates using JC-1, a small positively charged molecule that fluoresces green when dispersed and red when aggregated under ultraviolet (UV) illumination (Garner and Thomas, 1999). Gradient strength can then be estimated from the ratio of red aggregate fluorescence to green fluorescence of the dye monomer. Third, electron flow through the electron transport chain (ETC) provides the energy necessary to establish a proton gradient (Brand and Murphy, 1987), such that increased electron flow should produce a corresponding increase in mitochondrial membrane potential. When isolated mitochondria are incubated with the compound MTT (3- (4,5-dimethylthizol-2-yl) diphenyltetrazolium bromide), MTT accepts electrons from the ETC, forming a purple formazan product, the quantity of which positively correlates with electron flow (Liu *et al.,* 1997). Finally, because O_2_ is the final electron acceptor for the production of ATP (Chance and Williams, 1955), total O_2_ consumption represents an integrated measure of the ATP production capacity of an organism. Aquatic respirometry using a Clark-type electrode can detect changes in O_2_ concentrations over time, allowing us to evaluate the rate at which whole organisms produce ATP. Because ectotherms respire at higher rates at elevated temperatures, heat stress can reveal inefficiencies in ATP production (Heise *et al.,* 2003; Abele *et al.,* 2007), allowing us to also use aquatic respirometry to compare ATP production in ambient *vs.* stress-inducing temperatures.

We adapted these well-established mitochondrial functional assays, to date employed exclusively in model organisms, to probe the number of mtDNA copies relative to the nuclear genome, estimate the strength of the electrochemical gradient, quantify the flow of electrons through the electron transport chain (ETC), and to quantify and compare total organismal O_2_ consumption under heat stress across asexual *P. antipodarum* lineages. Together, these analyses revealed substantial variation for mitochondrial function across distinct asexual lineages at all three levels of biological organization. More broadly, the assays adapted here provide a suite of useful experimental tools for investigating mitochondrial function in *P. antipodarum* and, potentially, other mollusks.

## MATERIALS AND METHODS

### Snail husbandry

We compared mitochondrial function across a diverse array of asexual *P. antipodarum* lineages (Neiman *et al.,* 2011; Paczesniak *et al.,* 2013; Table S1) reared under identical conditions for multiple generations. Asexual lineages were chosen for functional assays to represent the range of mitochondrial genetic diversity found in New Zealand populations (as demonstrated by Neiman and Lively, 2004; Neiman *et al.,* 2011; Paczesniak *et al,* 2013) and to maximize our ability to compare across functional assays. Because there is notably high genetic diversity within asexual assemblages (Jokela *et al,* 2003; Paczesniak *et al,* 2013) and marked across-lake genetic structure (Neiman and Lively, 2004; Paczesniak *et al.,* 2013) in New Zealand *P. antipodarum,* and because nearly all of the lineages were from different lakes (Table S1), we can confidently interpret across-lineage variation in our various measures of mitochondrial function as representing genetic variation for these traits. Asexuality was established for each lineage by determining ploidy using flow cytometry (sexual *P. antipodarum* are diploid, asexuals are polyploid), as described in Neiman *et al.* (2011). We did not determine ploidy level for field-collected snails, which is why we did not use these individuals in lineage-level comparisons of mitochondrial performance. Adult female snails were selected arbitrarily from lineage populations or field collections for each assay. Following standard laboratory protocols for *P. antipodarum* (for example, Zachar and Neiman, 2013), snails were housed at 16°C on an 18 hr light/6 hr dark schedule and fed *Spirulina* algae 3x per week.

### Mitochondrial function at the genomic level

To test whether asexual *P. antipodarum* exhibited across-lineage phenotypic variation for mitochondrial copy number, we used quantitative PCR (qPCR) to estimate mtDNA copy number relative to a putatively single-copy nuclear gene in six asexual triploid lineages. To identify a suitable nuclear-encoded gene to use as a control, we performed an all-by-all BLAST search of the transcriptomes available at http://bioweb.biology.uiowa.edu/neiman/blastsearch.php (Wilton *et al.,* 2013) to identify assembled transcripts that hit themselves and only themselves. We randomly selected 30 transcripts satisfying that criterion, designed primers, amplified sequences *via* PCR, and sequenced each transcript on an ABI 3730 DNA Analyzer (Applied Biosystems, Foster City, CA). Because *rad21* was the most consistent performer in PCR amplification experiments, we used our sequencing information to design new internal primers designed to produce a 264-bp product (F: 5’-GATTCCAACAACTGATGTTTG −3’, R: 5’-CAAAACTTACTCTAAATCTGC−3’) for use as a nuclear genome standard in qPCR experiments. We then designed primers to produce a 194-bp amplicon from *cytB* (F: 5’- TATGAATATTCAGATTTTTTAAATA−3’, R: 5’- CCTTAACTCCTAATCTTGGTAC−3’), our mitochondrial standard. For measurement standards, each of these products was cloned from total DNA from a single *P. antipodarum* individual into the pGEM T-Easy plasmid vector (Promega Corp., Madison, WI). Linearized plasmids were diluted in the presence of carrier human genomic DNA to produce samples containing 300 - 300,000 copies of either the nuclear or mitochondrial amplicon. To evaluate mtDNA copy numbers, total DNA was isolated from three to five individual snails from each of the six lineages using a DNeasy Plant Mini Kit (Qiagen, Valencia, CA). We used this DNA to amplify nuclear and mitochondrial targets in triplicate in separate reactions on the same plate, together with serial dilutions of the cloned standards, using quantitative PCR on a StepOne Plus real-time thermal cycler (Applied Biosystems, Foster City, CA). We converted quantitation cycle values (Cq, the PCR cycle at which amplification products accumulated above a defined threshold) from snail samples into copy numbers using standard curves generated from the cloned standards, as in Miller *et al.* (2003). We then used this information to determine the ratio of mitochondrial to haploid nuclear genome copies for each sample. Finally, we compared inferred mtDNA copy number across lineages using a one-way ANOVA and pairwise t-tests, ensuring that mtDNA copy numbers were normally distributed (Shapiro-Wilks *W* = 0.96, *p* = 0.23) and that variances between lineages were not significantly different (Levene’s *F* = 0.63, *p* = 0.68). All statistical tests were performed in R v 3.2.4 (R Core Team, 2012), and all plots were produced using the *car* R package (Fox and Weisberg, 2011).

### Mitochondrial function at the organellar level

Except where noted, all reagents were obtained from Sigma-Aldrich, St. Louis, MO.

#### JC-1 assay

To test for genetic variation in mitochondrial function in *P. antipodarum* in terms of mitochondrial membrane potential, we assayed mitochondrial membrane potentials using the JC-1 dye assay in 8-10 individual adult female snails from six distinct asexual lineages representing a diverse subset of the natural mitochondrial haplotype diversity found in New Zealand (Neiman *et al.,* 2011; Paczesniak *et al.,* 2013; Table S1). We used the uncoupler carbonyl cyanide m-chlorophenyl hydrazone (CCCP) to abolish the electrochemical gradient across the mitochondrial inner membrane in replicate subsamples, allowing us to control for background levels of fluorescence of JC-1 and of mitochondrial membranes unrelated to mitochondrial function.

For each snail, we first removed its shell and briefly washed the collected tissues by centrifugation at 600 x g in extraction buffer (10.0 mM HEPES [pH 7.5], 0.2 M mannitol, 70.0 mM sucrose, 1.0 mM EGTA). We rapidly homogenized these tissues on ice in extraction buffer containing 2 mg/ml fatty acid-free bovine serum albumin (fafBSA) using a micropestle and then centrifuged the homogenate at 4°C for 5 minutes at 600 x g. The supernatant was recovered and held on ice separately while the pellet was re-homogenized and centrifuged again, as above. The pooled mitochondrial-enriched supernatant was centrifuged at 12,000 x g for 10 minutes and the pellet resuspended in buffer containing 10.0 mM HEPES (pH 7.5), 0.25 M sucrose, 1.0 mM ATP, 0.08 mM ADP, 5.0 mM sodium succinate, 2.0 mM K_2_HPO_4_, and 1mM DTT. We divided each sample into three subsamples of 30 μl and added 500 μl assay buffer containing 20.0 mM MOPS (pH 7.5), 110.0 mM KCl, 10mM ATP, 10.0 mM MgCl_2_, 10.0 mM sodium succinate, and 1.0 mM EGTA. The first subsample was incubated with buffer alone, to monitor background fluorescence, the second subsample was incubated with 2 μM JC-1 (Calbiochem, San Diego, CA), and the third with 2 μM JC-1 and 30 μM CCCP. All three subsamples were incubated in the dark at 37°C for 20 minutes, after which the ratio of red: green fluorescence was determined using flow cytometry (Becton Dickinson FacsCalibur, Franklin Lakes, NJ). Ungated data were collected from several hundred to several thousand mitochondrial particles per sample, on forward scatter (FSC) and side scatter (SSC), FL1 (green fluorescence, log scale), and FL2 (red fluorescence, log scale). We plotted FL1 *vs.* FL2 for each subsample after gating out debris in FlowJo v 10.0.8 (FlowJo, Ashland, OR), and derived the ratio of red to green for each particle. Unstained subsamples showed very low background autofluorescence that did not vary across lineages. Stained subsamples uncoupled by CCCP had low red:green ratios representative of dissipated charge gradients, while subsamples without uncoupler contained an additional fraction of mitochondrial particles with higher ratios, representing coupled mitochondria. We used one-way ANOVA with lineage as a random factor to compare the median red: green ratio of particles in this final fraction across lineages following log transformation of red: green fluorescence ratios (Shapiro-Wilks *W* = 0.858, *p* = 7.131 × 10^−6^ prior to log transformation; Shapiro-Wilks *W* = 0.969, *p* = 0.15 following log transformation). Our ANOVA approach required a White adjustment (MacKinnon and White, 1985) because ratios of red: green fluorescence exhibited unequal variances across lineages, (Levene’s *F* = 3.093, *p* = 0.016).

#### MTT assay

We next tested whether there exists genetic variation for electron flux through the ETC in *P. antipodarum* by comparing MTT reduction across asexual lineages. We resuspended mitochondrial pellets pooled from 3-4 snails (obtained as described above) in 100 μl buffer (125 mM KCl, 2 mM K_2_HPO_4_, 1 mM MgCl_2_, and 20 mM HEPES, adjusted to pH 7.4 with KOH) with 6 mM succinate as an energy substrate. We then added these resuspended mitochondria to wells in 96-well plates. The MTT reaction was initiated by adding 10 μl of 2.5 mg/ml MTT to each well. We then incubated the plate for 2 hours at 37°C to allow electrons from the ETC to reduce MTT. Next, a 20% SDS 50% dimethylformamide solubilization solution was applied to each well and the plate was incubated overnight, after which the reduced formazan product was measured as A_570_ in a XL-800 microplate reader (Bio-Tek Instruments, Winooski, VT). We determined background absorbance from duplicate mitochondrial samples incubated without an energy substrate and subtracted this background value from all readings. Phosphate-buffered saline was used as a negative control and 0.1 mM dithiothreitol as a positive control for MTT reduction. A fraction of each original mitochondrial sample was lysed in SDS and then used in a bicinchoninic acid assay (Smith *et al.,* 1985) to determine protein concentration. MTT reduction was expressed as A_570_/ug protein. We performed a log transformation of the raw MTT values (Shapiro-Wilks *W* = 0.840, *p* = 2.436 × 10^−5^) to meet the assumptions of normality (Shapiro-Wilks *W* = 0.961, *p* = 0.140) and test for unequal variances (Levene’s *F* = 0.964, *p* = 0.452). We then compared log-transformed MTT values using a one-way ANOVA with lineage as a random factor to test whether different asexual lineages exhibited different levels of electron flow through the ETC.

### Mitochondrial function at the organismal level

To test for variation in O_2_ consumption in response to heat stress, we first needed to establish the range of heat stress likely to alter mitochondrial function in *P. antipodarum.* We accomplished this goal by using a behavioral assay that measures the amount of time that a snail takes to right itself when placed ventral side-up to compare righting ability in 13 asexual lineages (N = 10 individuals per lineage) across three temperature treatments (16°C, 22°C, 30°C). Righting ability is a commonly used method to gauge levels of snail stress (for example,Orr *et al.,* 2007). For example, snails exposed to hypoxic conditions exhibit increased righting time (and elevated HSP70 gene expression) compared to unexposed snails (Fei *et al.,* 2008). We began by incubating adult female *P. antipodarum* in carbon-filtered tap water (that is, the water in which the snails are housed) at the test temperature for 1 hour. Next, we placed snails ventral side-up in a petri dish and measured the number of seconds elapsed until the snail righted itself, up to 180 seconds.

After determining that snails do appear to be stressed by elevated temperatures (see Results, Figure S1), we tested whether there exists genetic variation for O_2_ consumption under heat stress by performing closed-system aquatic respirometry on seven asexual lineages of *P. antipodarum* (N = 10 per lineage) at the same three incubation temperatures used for the righting assay (16°C, 22°C, 30°C) with a Strathkelvin Instruments RC200a respiration chamber, a 892 Oxygen Meter, and a 1302 Clark-type oxygen electrode (Strathkelvin Instruments, Motherwell, UK). We calibrated the electrode daily using the solubility of O_2_ at each respective temperature (16°C - 309.0 μmol/liter, 22°C - 279.0 μmol/liter, 30°C - 236.0 μmol/liter). A high calibration point was obtained by stirring carbon-filtered water vigorously for 30 minutes prior to n-calibration, while we used a 2% sodium sulfite solution as a low calibration standard. We incubated each snail at the prescribed temperature for one hour prior to measurement, placed snails into the cell chamber, and obtained O2 concentration readings for each snail every second for 1 hour. We maintained constant temperature in the respiration chamber by pumping temperature-controlled water into the cell chamber’s water jacket. Upon completion of the 1-hour test period, we blotted each snail dry and measured its wet mass on a Denver Instruments Cubis Analytical Balance (Denver Instrument, Bohemia, NY). After correcting for snail wet mass, we then used a two-way ANOVA to address whether the fixed factor of temperature, the random factor of lineage, and the interaction between temperature and lineage affected the dependent variable of mass-corrected O_2_ consumption.

### Comparison of mitochondrial functional assays

Our mitochondrial function assays were aimed at different elements of mitochondrial performance and different levels of biological organization; whether and to what extent these assays are measuring related *vs.* orthogonal determinants of mitochondrial function remains unclear, especially as newly applied to *P. antipodarum.* We addressed this question by performing all 15 possible pairwise comparisons of mitochondrial functional assays (that is, mtDNA copy number, mitochondrial membrane potential, electron flux, O_2_ consumption at 16°C, O_2_ consumption at 22°C, and O_2_ consumption at 30°C), comparing the mean trait value per lineage across assays using Spearman’s rank correlation (as implemented by the *Hmisc* R package [Harrell, 2008]) and correcting for multiple comparisons using the Holm procedure (Holm 1979). For these analyses, we included two additional sample populations that were not used in any of the tests for across-lineage variation: one field-collected sample from a lake with a high frequency of sexual individuals (“KnSF12”) and an inbred diploid sexual lineage that has been maintained in the lab for >20 generations (“Y2”, Table S1).

## RESULTS

### Mitochondrial function at the genomic level

To test whether *P. antipodarum* exhibits heritable variation for mtDNA copy number, we compared qPCR amplification of the mitochondrially encoded *cytB* locus to amplification of a putatively single copy nuclear gene, *rad21,* in six asexual lineages of *P. antipodarum* reared in a common-garden setting (N = 3-8 per lineage). The mean (+/- SD) number of *cytb* copies to the number of *rad21* copies was 13.72 (+/- 3.57), though we also found significant differences in this ratio across asexual lineages (one-way ANOVA, *F_5_*, _25_ = 2.72, *p* = 0.043, Figure 1), indicating that lineages differ in mtDNA copy number. We performed pairwise t-tests of mtDNA copy number ratio among each pair of lineages, which revealed that a single lineage (Gr5; mean copy number ratio [+/-SD] = 9.42 [+/−1.48]) with very low copy number appeared to be driving the significant among-lineage variance. This conclusion is supported by the fact that when Gr5 is excluded, other asexual lineages exhibit no differences in copy number (one-way ANOVA, F_4, 21_ = 0.55, *p* = 0.70).

**Figure 1.**
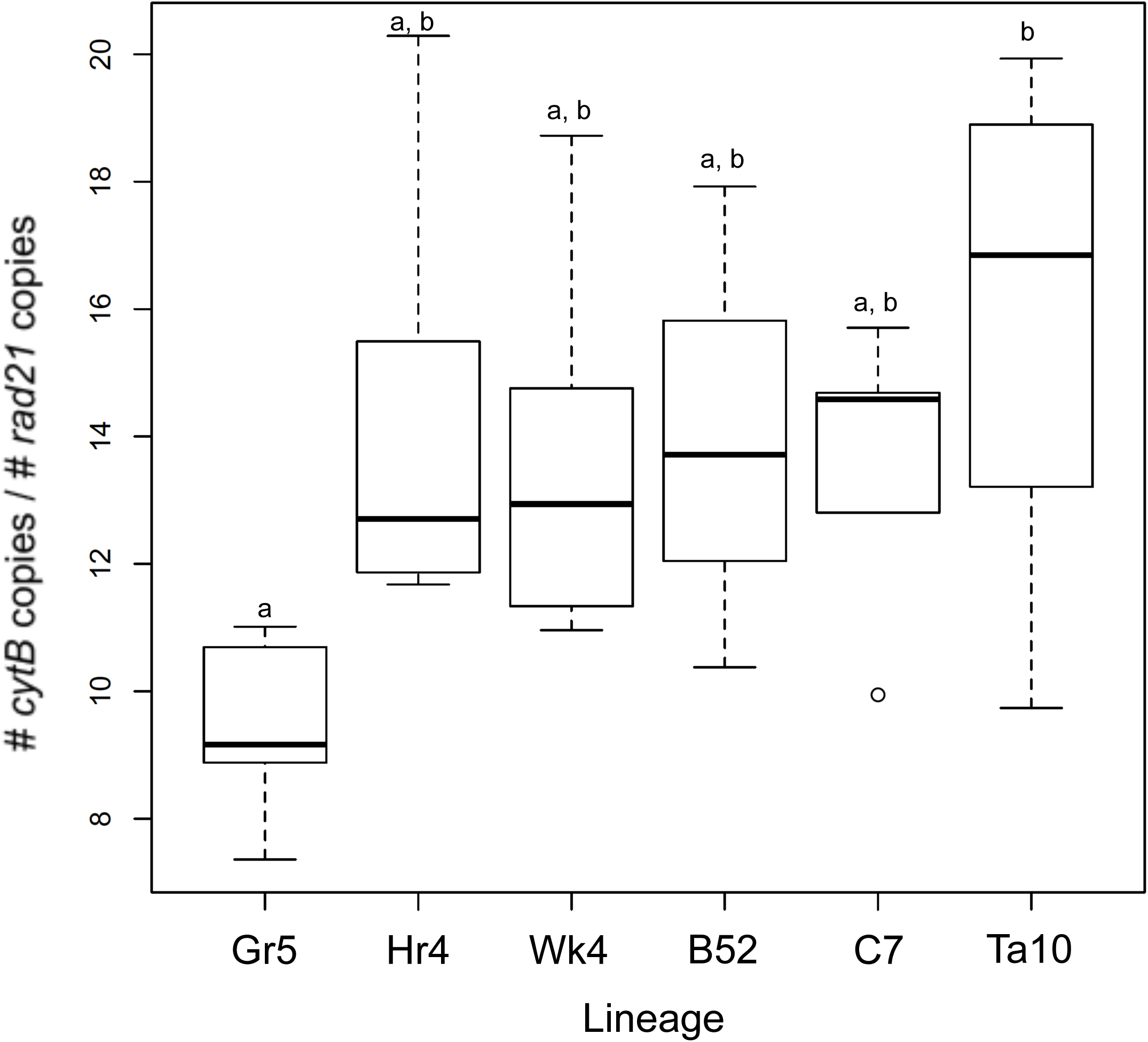
mtDNA copy number variation in six asexual lineages of *P. antipodarum.* Box-and-whisker plot depicting qPCR estimates (rank ordered by median) of *cytB* copy number relative to a putatively single-copy nuclear gene, *rad21.* Boxes represent Inner Quartile Ranges (IQR), with error bars extending 1.5x beyond IQRs. Data points falling outside whiskers are denoted by open circles. Shared lowercase letters indicate *p* > 0.05 for pairwise t-tests corrected for multiple comparisons using the Holm procedure for multiple comparisons. N = 10 individuals for all lineages.

### Mitochondrial function at the organellar level

#### >JC-1

We compared membrane potential among mitochondria isolated from six asexual lineages of *P. antipodarum* using the JC-1 assay, in which the ratio of red to green fluorescence indicates the relative strength of the proton gradient. Our comparisons revealed significant differences in log-transformed ratios of red: green across asexual lineages (Welch’s one-way ANOVA, F_5, 52_ = 6.628, *p* = 7.671 × 10^−5^, Figure 2a). We next performed *posthoc* pairwise comparisons between lineages using Welch’s t-tests (to reflect unequal variances) and the Holm procedure for Bonferroni correction (Holm 1979), which revealed three significantly different groups (*p <* 0.05 after correcting for multiple comparisons) amongst the six lineages (Figure 2a). Unlike mtDNA copy number, this variation in mitochondrial membrane potential does not appear to be driven by a single lineage. Because CCCP-uncoupled samples did not vary across lineages (one-way ANOVA, *F_5_*, _52_ = 1.37, *p* = 0.25), the variation observed for red: green fluorescence across lineages likely reflects true variation in mitochondrial membrane potential. In particular, these data suggest that under the current rearing and assay conditions, the Gr5 lineage appears to have the strongest mitochondrial membrane potential (mean [+/- SD] = 3.86 [+/- 1.55]) and the DenA lineage (mean [+/- SD] = 1.78 [+/- 0.47]) appears to have the weakest mitochondrial membrane potential.

**Figure 2.**
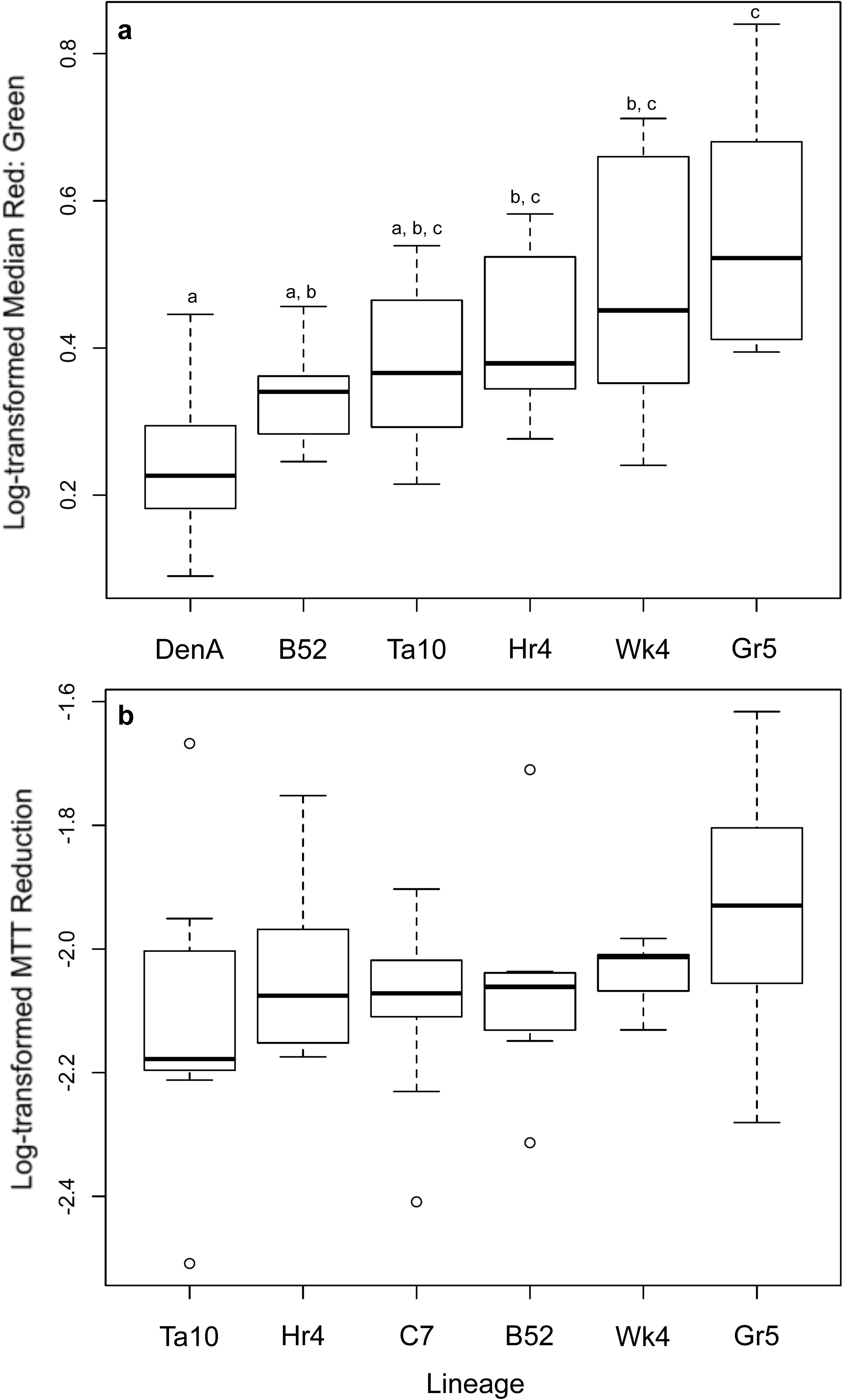
Organellar function of mitochondrial fractions isolated from six lineages of *P. antipodarum.* a) Estimate of mitochondrial membrane potential using the ratio of red to green JC-1 dye fluorescence from individual snails (N = 10 individuals for all lineages). Lineages were rank ordered by median with boxes representing IQRs; error bars extend 1.5x beyond IQRs. Data points falling outside error bars are denoted by open circles. Shared lowercase letters indicate *p* > 0.05 for pairwise Welch’s T-tests, corrected using the Holm procedure for multiple comparisons. b) Estimate of electron flux through OXPHOS pathway using the colorimetric MTT reduction assay from pooled mitochondrial extractions (3 snails pooled per replicate, 3 – 6 replicates per lineage). Lineages were rank ordered by median value with boxes representing IQRs and error bars extending 1.5x beyond IQRs. Data points falling outside error bars are denoted by open circles. Lineages did not appear to differ in MTT reduction.

#### MTT

We next compared electron flux through the ETC using the colorimetry-based MTT assay. We did not detect any differences amongst asexual lineages in MTT reduction (one-way ANOVA, *F*_5, 38_ = 0.75, *p* = 0.59, Figure 2b) despite relatively high statistical power for this comparison (= 0.92, as estimated by the *pwr* package in R [Champely, 2012]). This result therefore indicates that *P. antipodarum* lacks substantial genetic variation for electron flux though the ETC, at least under the benign environmental conditions in which we performed this assay.

### Mitochondrial function at the organismal level

Organismal O_2_ consumption under stressful conditions is expected to reflect electron acceptor turnover under maximal ATP production (Abele *et al.,* 2007), which is the logical basis for our application of heat stress to detect genetic variation in O_2_ consumption. We began by comparing righting time across ambient (16°C) and elevated (22°C, 30°C) temperatures and lineages with a Kruskal-Wallis test, and found that both factors significantly affected righting time (temperature: χ^2^ = 14.218, df = 2, *p* = 0.00082; lineage: χ^2^ = 122.64, df = 12*,p* < 2.2 × 10^−16^). *Post-hoc* Mann-Whitney U tests revealed that righting took ~ 37% longer at 30°C (mean [+/- SD] = 83.02 [+/- 73.74] seconds) than at 16°C (Mann-Whitney U = 8192, *p* = 0.025) and ~85% longer at 30°C than at 22°C (Mann-Whitney, *U* = 5996.5, *p* = 0.00014, Figure S1). While righting time was ~24% faster at 22°C (mean [+/- SD] = 48.82 [+/- 56.09] seconds) than at 16°C (mean [+/- SD] = 62.50 [+/- 63.26] seconds), this difference was not significant (Mann-Whitney, *U* = 9244.5, *p =* 0.075). These results are consistent with other righting time assays from other snail species in that some amount of stress decreases righting time (for example, presence of predators,Orr *et al.,* 2007), and some stimuli increase righting time (for example, hypoxia,Fei *et al.,* 2008), suggesting that our observed responses indicate stress in *P. antipodarum* exposed to elevated temperature (here, 30°C).

To test for genetic variation in O_2_ consumption in heat-stressed *P. antipodarum,* we performed closed-system aquatic respirometry for seven asexual lineages (N = 10 per lineage) at 16°C, 22°C, and 30°C. Snail wet mass was significantly and positively correlated with O_2_ consumption (Spearman’s rho = 0.19, *p* = 0.0086, Figure S2). Because there is significant variation for snail wet mass across asexual lineages (Kruskal-Wallis, χ^2^ = 75.09, df = 6, *p* < 0.00010), we calculated the residuals of wet mass vs. O_2_ consumption using a linear model. Cube root-transformed, mass-corrected O_2_ consumption residuals were not significantly different from a normal distribution (Shapiro-Wilks *W* = 0.99, *p* = 0.060), allowing us to implement a linear regression model to test whether temperature and/or lineage affected O_2_ consumption. We found that elevated temperatures significantly affected mass-corrected O_2_ consumption (two-way

ANOVA, F_2, 175_ **=** 46.22, *p* = 2.2 x 10−^16^) and that there was a significant interaction between lineage and temperature (Figure 3, two-way ANOVA, F6, 175 = 3.40, *p* = 1.7 x 10^−4^). We also found a trend towards an effect of lineage on mass-corrected O2 consumption (two-way ANOVA, F_6_, _175_ = 2.12, *p* = 0.053). A series of one-way ANOVAs within each temperature treatment revealed that lineage had a significant effect on O_2_ consumption at 22°C (one-way ANOVA, F_6, 54_ = 2.83, *p* = 0.018) and at 30°C (one-way ANOVA, F_6, 62_ = 3.85, *p* = 0.0025), but not at 16°C (one-way ANOVA, F_6, 59_ = 2.10, *p* = 0.067). In particular, lineages responded differently to elevated temperature, with some lineages (for example, Ta10) exhibiting relatively high O_2_ consumption at high temperatures and others maintaining similar levels of O_2_ consumption (for example, Gn5) across temperatures (Figure 3). This result demonstrates that genetically distinct lineages of *P. antipodarum* consume different amounts of O_2_ in response to elevated temperature (22°C) and heat stress (30°C).

**Figure 3.**
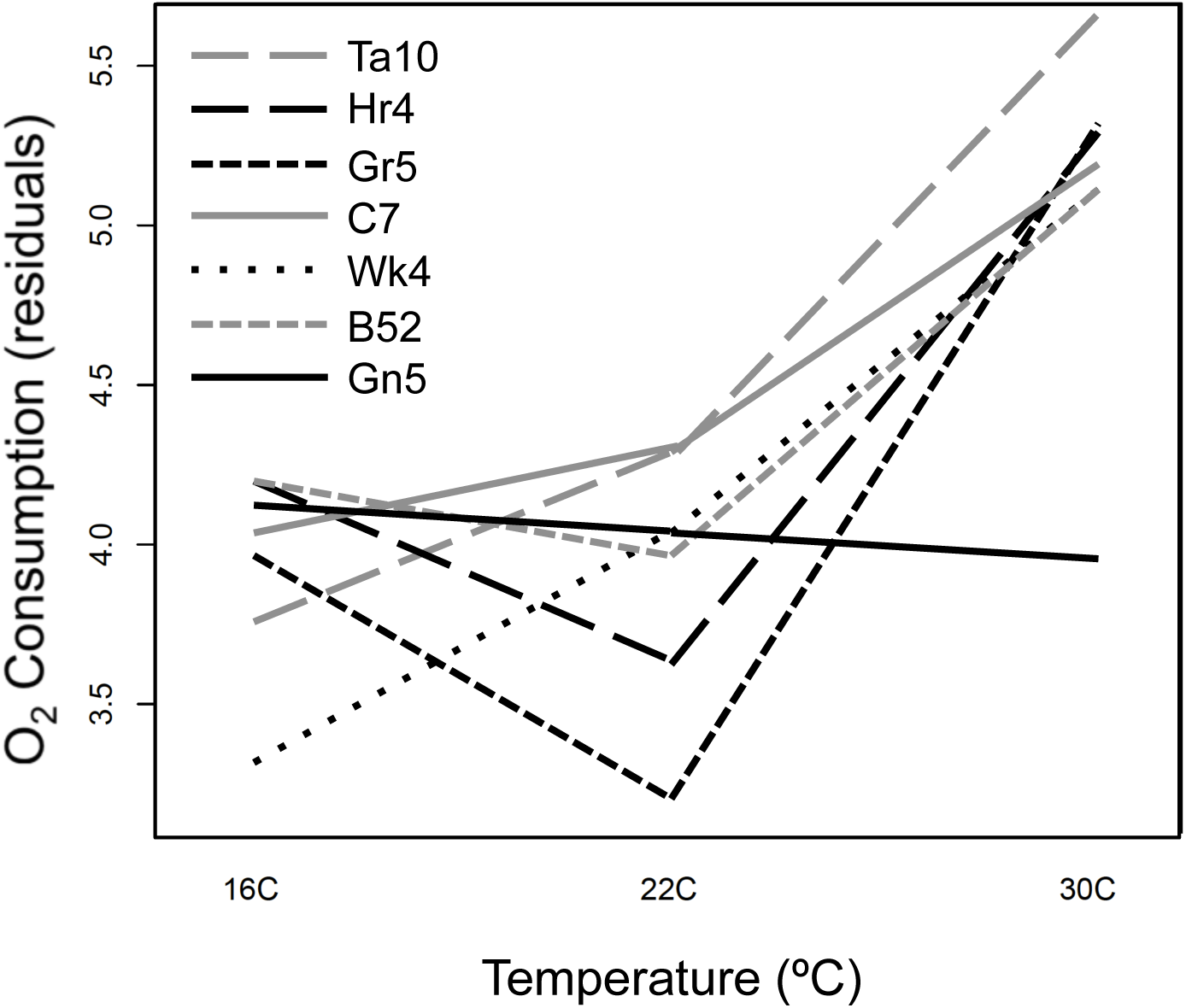
Interaction plot depicting relationship between O_2_ consumption residuals, temperature, and snail lineage. Lines indicate best-fit linear regression of O_2_ consumption across temperature pairs (for example, 16°C – 22°C) for seven asexual lineages of *P. antipodarum.* O_2_ consumption was measured for 10 individual snails at each temperature for each lineage.

### Comparison of mitochondrial functional assays

To determine whether and to what extent our different measures of mitochondrial function are associated, we performed Spearman’s rank correlation for each of the 15 possible pairwise comparisons of the assays for mtDNA copy number, log-transformed JC-1 red: green ratios, log-transformed MTT values, and cube root-transformed, mass-corrected O_2_ consumption residuals at each of our three study temperatures. None of the correlations remained significant after the Holm correction for multiple comparisons (Figure 4). The absence of significant relationships between assay results is not surprising in light of the fact that we had power of ~ 0.35 - 0.65 to detect a relationship between two variables at the *p* < 0.003 level, the alpha required by the Holm procedure. Power notwithstanding, there were some interesting potential trends that deserve further scrutiny. First, mtDNA copy number appears to be positively correlated with O_2_ consumption at 30°C among the six asexual lineages assayed (Spearman’s rho = 0.94, *p* = 0.0048, Figure 4e). Second, while the MTT assay required pooling of mitochondria from 3-4 individual snails per measurement and is thus somewhat less sensitive than the JC-1 assay, which only requires mitochondria from one snail, we did find a trend towards a positive correlation between electron flux and mitochondrial membrane potential in the six asexual and two sexual *P. antipodarum* lineages examined (Spearman’s rho = 0.81*,p* = 0.015, Figure 4f). Third, there electron flux appeared to be negatively correlated with O_2_ consumption at 22°C for the six asexual lineages (Spearman’s rho = −0.89, *p* = 0.019, Figure 4k). These tentative relationships between mtDNA, electron flux through the ETC, mitochondrial membrane potential, and organismal O_2_ consumption suggest that 1) the methods we employed here assay distinct yet associated mitochondrial phenotypes, and 2), evaluating mitochondrial performance at multiple levels of biological organization is necessary to describe adequately the phenotypic variation for mitochondrial function in *P. antipodarum.*

**Figure 4.**
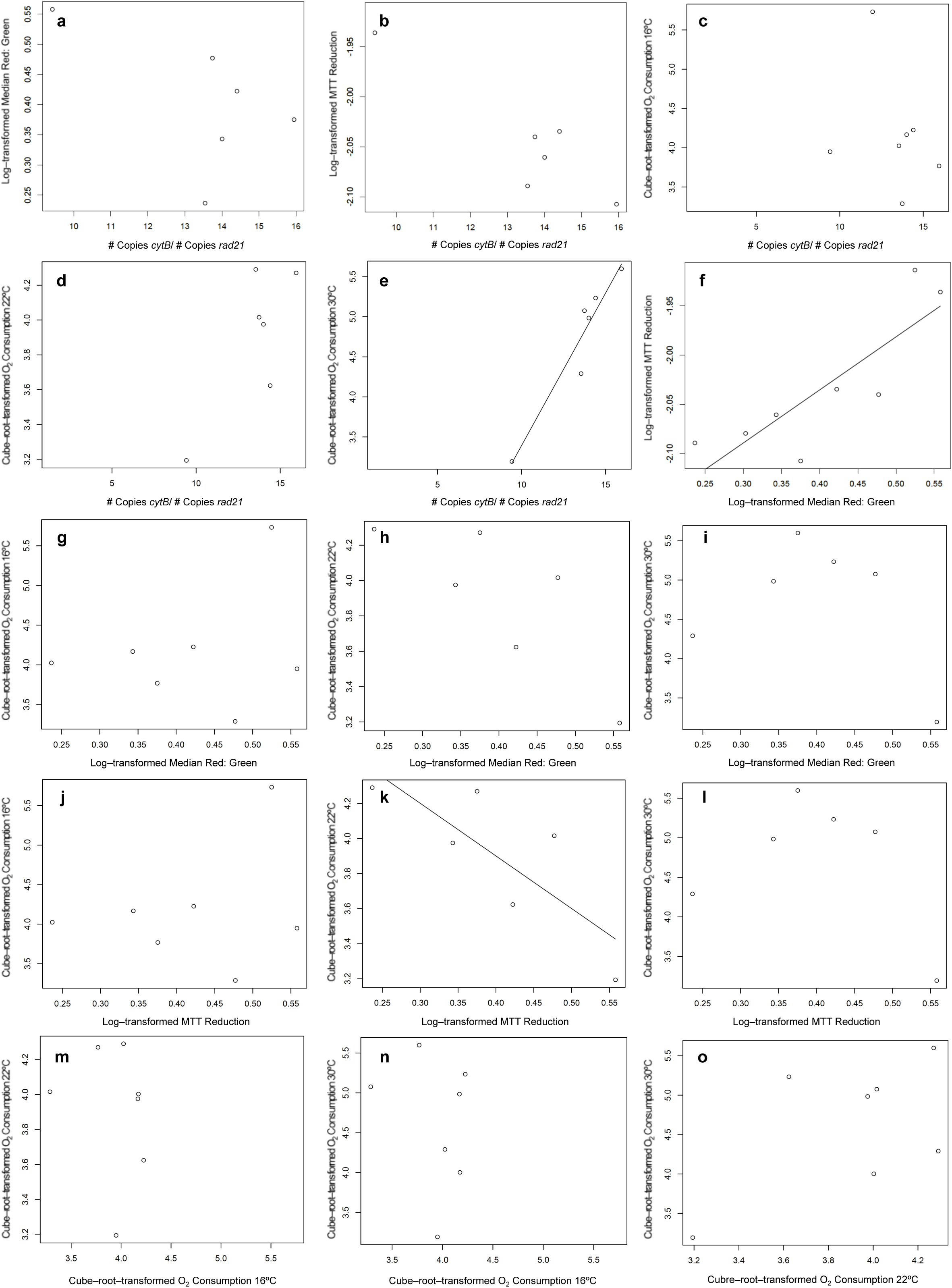
Comparisons of mitochondrial functional assays. Spearman’s rank correlations for all pairwise comparisons of mitochondrial functional assays. Best-fit linear regression lines (black) are only shown for those comparisons that were significant at the *p* < 0.05 level. a) mtDNA copy number vs. JC-1 (Spearman’s rho = −0.1, *p* = 0.82). b) mtDNA copy number vs. MTT reduction (Spearman’s rho = −0.33, *p* = 0.42). c) mtDNA copy number vs. O_2_ consumption at 16°C (Spearman’s rho = −0.14, *p* = 0.76). d) mtDNA copy number vs. O_2_ consumption at 22°C (Spearman’s rho = 0.2, *p* = 0.70). e) mtDNA copy number vs. O_2_ consumption at 30°C (Spearman’s rho = 0.94, *p* = 0.0048). f) JC-1 vs. MTT reduction (Spearman’s rho = 0.81, *p* = 0.015). g) JC-1 vs. O_2_ consumption at 16°C (Spearman’s rho = 0.0, *p* = 1.0). h) JC-1 vs. O_2_ consumption at 22°C (Spearman’s rho = −0.71, *p* =0.11). i) JC-1 assay vs. O_2_ consumption at 30°C (Spearman’s rho = −0.09, *p* = 0.87). j) MTT reduction vs. O_2_ consumption at 16°C (Spearman’s rho = 0.5, *p* = 0.25). k) MTT reduction vs. O_2_ consumption at 22°C (Spearman’s rho = −0.89, *p* = 0.019). l) MTT reduction vs. O_2_ consumption at 30°C (Spearman’s rho = −0.43, *p* = 0.40). m) O_2_ consumption at 16°C vs. O_2_ consumption at 22°C (Spearman’s rho = −0.39, *p* = 0.38). n) O_2_ consumption at 16°C vs. O_2_ consumption at 30°C (Spearman’s rho = −0.14, *p* = 0.76). o) O_2_ consumption at 22°C vs. O_2_ consumption at 30°C (Spearman’s rho = 0.32, *p* = 0.48).

## DISCUSSION

### Genetic variation for mitochondrial function in asexual lineages

We used obligately asexual lineages of *Potamopyrgus antipodarum,* a freshwater New Zealand snail, to test for genetic variation in mitochondrial function in a common-garden setting. We found significant levels of variation at all three levels of biological organization that we assayed, 1) mtDNA copy number, 2) mitochondrial membrane potential, and 3) variation in O_2_ consumption in response to heat stress. These results demonstrate that substantial variation for mitochondrial function exists across asexual lineages of this species.

While the extent to which the variation in mitochondrial function described here contributes to fitness in *P. antipodarum* remains to be directly evaluated, the close link between mitochondrial function and fitness in other organisms (Chen *et al.,* 2007; Dowling, 2014) suggests that phenotypic variation across mitonuclear genotypes could have major implications for asexual lineage success. In particular, asexual *P. antipodarum* are known to harbor high mtDNA mutational loads relative to sexual lineages (Neiman *et al.,* 2010; Sharbrough *et al.,* 2016), and heat stress response can affect fecundity (Dybdahl and Kane, 2005) and respiration rates (Hudson, 1983, present study) in *P. antipodarum.* These findings suggest that variation in mitochondrial function might very well confer fitness consequences, especially amongst asexual lineages that are experiencing stressful conditions.

The substantial across-lineage variation that we discovered provides functional evidence of high levels of asexual phenotypic diversity in *P. antipodarum,* consistent with previous reports that asexual *P. antipodarum* harbor substantial genetic (Jokela *et al.,* 2003; Paczesniak *et al.,* 2013) and phenotypic (Neiman *et al.,* 2009, 2013; Kistner and Dybdahl, 2013) diversity. The multiple separate transitions to asexuality in *P. antipodarum* (Neiman *et al.,* 2011; Paczesniak *et al.,* 2013) may help explain the substantial levels of variation found here, as asexual lineages represent “snapshots” of local sexual population diversity (Dybdahl and Lively, 1995; Jokela *et al.,* 1997).

The mitochondrial phenotypes we observed in one lineage, Gr5, were particularly distinct: snails from this lineage exhibited relatively low mitochondrial copy number and high mitochondrial membrane potential and electron flow, as well as relatively low O_2_ consumption at 22°C. Together, these phenotypic values suggest that the Gr5 lineage exhibits relatively high mitochondrial function, indicating that genetic dissection of its mitochondrial haplotype may prove illuminating. Further comparisons with other *P. antipodarum* lineages and in other conditions will provide substantial insight into the relative fitness of this particular mitonuclear combination. Future studies should also focus on a particular mitochondrial haplotype that appears to be especially common amongst asexual *P. antipodarum* (Neiman *et al.,* 2011; Paczesniak *et al.,* 2013). Paczesniak *et al.* (2013)showed that this haplotype is often found in divergent nuclear backgrounds, consistent with a scenario in which this haplotype is spreading into new populations and lineages. Evaluating mitochondrial function of the common mitochondrial haplotype against a variety of nuclear backgrounds and in various biologically relevant conditions would shed light on intraspecific mitonuclear coevolution and whether asexuality contributes to decreased mitochondrial function in *P. antipodarum*, as the mutational hypotheses for sex would predict.

### Relationships among mitochondrial functional assays

All else being equal, stronger mitochondrial membrane potentials and greater electron flow should indicate relatively high mitochondrial performance. The relationships between whole-organismal O_2_ consumption and mitochondrial performance or between mtDNA copy number and mitochondrial performance are expected to be more complex and to reflect compensatory mechanisms at the cellular and/or organismal level. Because other factors (for example, environmental conditions, local adaptation) are virtually certain to influence mitochondrial function, it is also possible that high mitochondrial performance in common-garden conditions does not reflect mitochondrial performance in nature. Despite this caveat, the tools developed here will provide an important starting point with which to interrogate mitochondrial function in non-model systems.

Although the number of asexual lineages included in the present study was relatively small, we did observe some interesting tentative relationships between mtDNA copy number, electron flux through the ETC, mitochondrial membrane potential, and organismal O_2_ consumption (Figure 4). The observation that lineages with high mtDNA copy numbers consume more O_2_ at elevated temperatures than lineages with lower mtDNA copy numbers may reflect saturation of OXPHOS pathways during heat stress in lineages with low mtDNA copy numbers. Compensation for reduced mitochondrial function by increasing mtDNA copy number has been observed in human tissues carrying a variety of small deletion mutations (Bai and Wong, 2005), although the relationship between mtDNA copy number and respiratory capacity remains complex (Moraes, 2001; Montier *et al,* 2009). The positive relationship between electron flux and mitochondrial membrane potential is not particularly surprising in light of the fact that as electrons pass through the ETC, a corresponding increase in mitochondrial membrane potential is expected as H_+_ ions are released into the intermembrane space (Chen, 1988). The association between relatively low electron flux and high O_2_ consumption is more difficult to understand because O_2_ consumption is expected to increase with electron flux (Jastroch *et al.,* 2010). In general, relationships between organismal-level O_2_ consumption and organelle-level electron flux are difficult to disentangle because many layers of respiratory regulation can contribute to increased organismal O_2_ consumption beyond ETC inefficiencies (Brand and Nicholls, 2011). A particularly relevant example is provided by combined observations that electrons leaking back across the inner membrane through uncoupling proteins contribute to electron flux but not O_2_ consumption (reviewed in Jastroch *et al,* 2010) and that elevated temperatures in a marine mollusk have been shown to increase electron leakage of this type (Abele *et al,* 2002), such that connecting organismal O_2_ consumption to organellar electron flux is a non-trivial exercise. Understanding the relationship between organellar and organismal variation in mitochondrial function will provide helpful context for interpreting functional variation in *P. antipodarum,* and future efforts towards examining these relationships in more detail are necessary. More broadly, the methods described here can be easily adapted to other non-model organisms, especially mollusks, providing a new means of quantifying the genotype-phenotype relationships of mitonuclear interactions.

## DATA ARCHIVING

All data will be made available on Dryad upon acceptance.

## ACKNOWLEDGEMENTS

We thank JD Woodell for assistance with aquatic respirometry and Claire Tucci, Meagan Luse, and Nikhil Puttagunta for their help measuring snail righting behavior. We acknowledge several anonymous reviewers for helpful comments on previous iterations of this manuscript.

## SUPPLEMENTARY MATERIAL TITLES TO FIGURES AND LEGENDS

**Figure S1. Interaction plot depicting relationship between temperature, lineage, and righting time in *P. antipodarum.*** Lines represent best-fit lines between righting times for a given lineage across 16°C - 22°C and 22°C - 30°C. Red lines represent asexual lineages; blue lines represent sexual lineages; black lines represent “mixed” field-collected populations. Snails righted themselves significantly more slowly at 30°C than 16°C (Mann-Whitney *U* = 8192, *p* = 0.025) and 22°C (Mann-Whitney, *U* = 5996.5, *p* = 0.00014), indicating that 30°C water temperatures represent a significant source of heat stress.

**Figure S2. Relationship between snail wet mass and oxygen consumption in *P. antipodarum.*** There was a significant positive correlation between snail wet mass and O_2_ consumption (Spearman’s rho = 0.19, *p* = 0.0086).

